# *Tsc1* haploinsufficiency leads to *Pax2* dysregulation in the developing murine cerebellum

**DOI:** 10.1101/2021.12.08.471768

**Authors:** Ines Serra, Ana Stravs, Catarina Osório, Maria Roa Oyaga, Martijn Schonewille, Christian Tudorache, Aleksandra Badura

## Abstract

Tuberous sclerosis complex 1 (*TSC1*) is a tumour suppressor gene that inhibits the mechanistic target of rapamycin (mTOR) pathway. Mutations in *TSC1* lead to a rare complex disorder of the same name, in which up to 50% of patients present with autism spectrum disorder (ASD). ASD is a highly prevalent, early-onset neurodevelopmental disorder, characterized by social deficits and repetitive behaviours, although the type and severity of symptoms show wide variability across individuals. Amongst different brain areas proposed to play a role in the development of ASD, the cerebellum is commonly reported to be altered, and cerebellar-specific deletion of *Tsc1* in mice is sufficient to induce an ASD-like phenotype. Given that the mTOR pathway is crucial for proper cell replication and migration, this suggests that dysregulation of this pathway, particularly during critical phases of cerebellar development, could contribute to the establishment of ASD.

Here, we used a mouse model of TSC to investigate gene and protein expression during embryonic and early postnatal periods of cerebellar development. We found that, at E18 and P7, mRNA levels of the cerebellar inhibitory interneuron marker *Pax2* were dysregulated. This was accompanied by changes in the expression of mTOR pathway-related genes and downstream phosphorylation of S6. Differential gene correlation analysis revealed dynamic changes in correlated gene pairs across development, with an overall loss of correlation between mTOR- and cerebellar-related genes in *Tsc1* mutants compared to controls. We corroborated the genetic findings by characterizing the mTOR pathway and cerebellar development on protein and cellular levels with Western blot and immunohistochemistry. We found that Pax2-expressing cells were hypertrophic at E18 while, at P7, their number was increased and maturation into parvalbumin-expressing cells delayed. Our findings indicate that E18 and P7 are crucial time points in cerebellar development in mice that are particularly susceptible to mTOR pathway dysregulation.

**Manuscript contribution to the field:** ASD is one of the most prevalent neurodevelopmental disorders, however little is known about the shared mechanisms underlying its aetiology. At the anatomical level, the cerebellum has been identified as one of the key structures involved in the development of ASD, whereas at the molecular level, mutations in the mTOR signalling pathway, essential for cell growth and proliferation, carry a high genetic risk for this disorder. We used a haploinsufficient tuberous sclerosis complex 1 (*Tsc1*) mouse model to investigate the effects of mTOR overactivation in the developing cerebellum. *Tsc1* inhibits the mTOR pathway, and mice with cerebellar-specific deletion of *Tsc1* have been shown to harbour an ASD-like phenotype. We found that Pax2 expression in the cerebellum is dysregulated at prenatal and early postnatal time points, leading to a delayed maturation of inhibitory interneurons. Our findings indicate that mTOR overactivity in the cerebellum selectively affects the development of cerebellar interneurons. This finding is in line with other studies, which found decreased numbers of inhibitory interneurons in other models of ASD. Therefore, deficits in the maturation of the inhibitory signalling could be one of the mechanisms integrating high-risk mutations that underlie ASD aetiology.

## Introduction

The mechanistic target of rapamycin (mTOR) pathway is a highly complex, conserved and ubiquitous signalling avenue involved in biomass synthesis, growth and cell proliferation (1). Specifically during brain development, it has been proposed that a tight regulation of mTOR signalling is required for sustaining cell cycle length and re-entry, defining pluripotency status and triggering differentiation (2–4). However, how the mTOR pathway affects distinct lineages of differentiating cells is still largely unexplored. Supporting a crucial role for the mTOR signalling in brain development, mutations along this pathway frequently lead to complex monogenic neurodevelopmental disorders (also known as *mTORopathies*), characterized by heterogeneous neuropsychiatric phenotypes that include megalencephaly, epilepsy, intellectual disability and autism spectrum disorder (ASD) (5,6).

The prototypical mTORopathy is tuberous sclerosis complex (TSC), a rare autosomal dominant disorder affecting 1 in 6000 people, that arises from heterozygous mutations in the *TSC1* or *TSC2* genes (7,8). As TSC1 and 2, together with TBC1D7, form a tumour suppressor complex upstream of mTOR, loss of function of this complex leads to mTOR pathway overactivity (9,10). mTOR can be organized in two complexes, mTORC1 and mTORC2, characterized, among others, by the presence of Raptor and Rictor, respectively (11,12). While mTORC1 is primarily associated with growth and proliferation, mTORC2 regulates cytoskeleton organization and cell motility (13,14). Nonetheless, crosstalk between mTORC1 and mTORC2 is vast, and changes in the function of both complexes due to mTOR dysfunction were shown to alter dendritic arbour morphology and synaptic transmission (15). On the whole, the effects of mTOR overactivity in TSC patients lead to a multi-system phenotype that includes widespread hamartoma growth, high prevalence of epilepsy, and, in up to 50% of the patients, ASD (16,17).

ASD is characterized by deficits in social communication and interaction, and by the presence of restricted, repetitive, and inflexible behaviours (18). The World Health Organization (WHO) estimates that 1 in 160 children worldwide will present with ASD, although its prevalence is known to vary across nations (19). Despite this high prevalence, little is known about the molecular mechanisms that underlie ASD. This is due to a significant knowledge gap, particularly with respect to brain development, when even limited signalling alterations can translate into considerable brain function, connectivity and structural deficits (20–22). While there is no single major anatomical abnormality evident in all people with ASD, the cerebellum is a brain structure that has emerged as a significant putative contributor to the development of ASD phenotypes. In humans, damage to the cerebellum is the second largest factor contributing to the risk of developing ASD (23–25), while cerebello-cortical connectivity is often found to be impaired in people with ASD (26,27). In recent years, several studies showed that murine models with cerebellar-specific deletion or inactivation of genes affecting the mTOR pathway, replicate these human phenotypes, presenting with decreased social interaction, increased repetitive behaviours and inflexible learning (28–31). Together with the fact that many mTOR pathway genes are found to be enriched in the cerebellum (32,33), this suggests that this brain area may be particularly sensitive to changes in mTOR pathway regulation.

Here, we used a haploinsufficient *Tsc1* mouse model, mimicking the human genotype of TSC, to investigate the effects of TSC1 deficiency in the developing cerebellum. We found that genetic dysregulation of the mTOR pathway can be detected from E18, suggesting a compensatory down-regulation in response to the hyperactivity of this pathway. Changes to cerebellar development can also be found at this age and postnatally at P7. Specifically, we found that Pax2 expression at these time points is altered, indicative of a delay in its expression in *Tsc1^+/-^* mice. This culminated in slowed maturation and reduced parvalbumin expression. Overall, our data suggest that mTOR overactivity in the cerebellum preferentially affects the development of cerebellar interneurons, which could potentially promote the development of altered circuitry and, consequently, lead to behavioural deficits.

## Materials and Methods

### Mouse procedures

Timed pregnancies were established between wild-type C57BL/6 females (*Tsc1^+/+^*) (Charles River Laboratories) and *Tsc1*^tm1Djk^ (*Tsc1^+/-^*) males to obtain mixed *Tsc1^+/+^* and *Tsc1^+/-^* litters (34). Vaginal plugs were checked daily, and embryonic day 0 (E0) was defined when a plug was observed. Confirmed pregnant dams were individually housed. Mice were maintained on a standard 12h light/dark cycle, with access to food and water *ad libitum*.

For the collection of embryonic samples, pregnant dams were briefly anesthetized prior to cervical dislocation, and E15 (n = 8 mice per genotype) and E18 (n = 6 mice per genotype) embryos collected onto cold PBS on ice. For the collection of neonatal samples, P1 (n = 8 mice per genotype) and P7 (n = 8 mice per genotype) mice were anesthetized prior to decapitation.

Cerebellar tissue was dissected in cold PBS under a Zeiss Stemi SV6 Stereo microscope. qPCR and western blot samples were collected into TRI Reagent^®^ (T9424, Sigma) or dry ice, respectively, and kept at −80°C until used.

All experimental animal procedures were approved *a priori* by an independent animal ethical committee (DEC-Consult, Soest, The Netherlands), as required by Dutch law, and conform to the relevant institutional regulations of the Erasmus MC and Dutch legislation on animal experimentation.

### Real-time qPCR

#### Primer design

Seven genes of interest along the TSC-mTOR pathway (*Tsc1, Tsc2, Rictor, Rptor, Mtor, Rps6kb1* and *Rps6*) (1) and 5 genes representing distinct cerebellar lineages (*Pax2, Pax6, Calb1, Slc1a3* and *Gdf10*) (35) were targeted. Housekeeping genes were selected based on previous literature using embryonic and neonatal mouse brain tissue (36–38). Two housekeeping genes were selected per age: *Ywhaz* and *Sdha* were used for the E15 group, *Gusb* and *Sdha* for E18, and *Gusb* and *Ywhaz* for P1 and P7.

Primer pairs were adapted from literature or designed using *Primer-BLAST* (ncbi.nlm.nih.gov/tools/primer-blast) and *Ensembl* (m.ensembl.org) (Table 1).

**Table 1:**
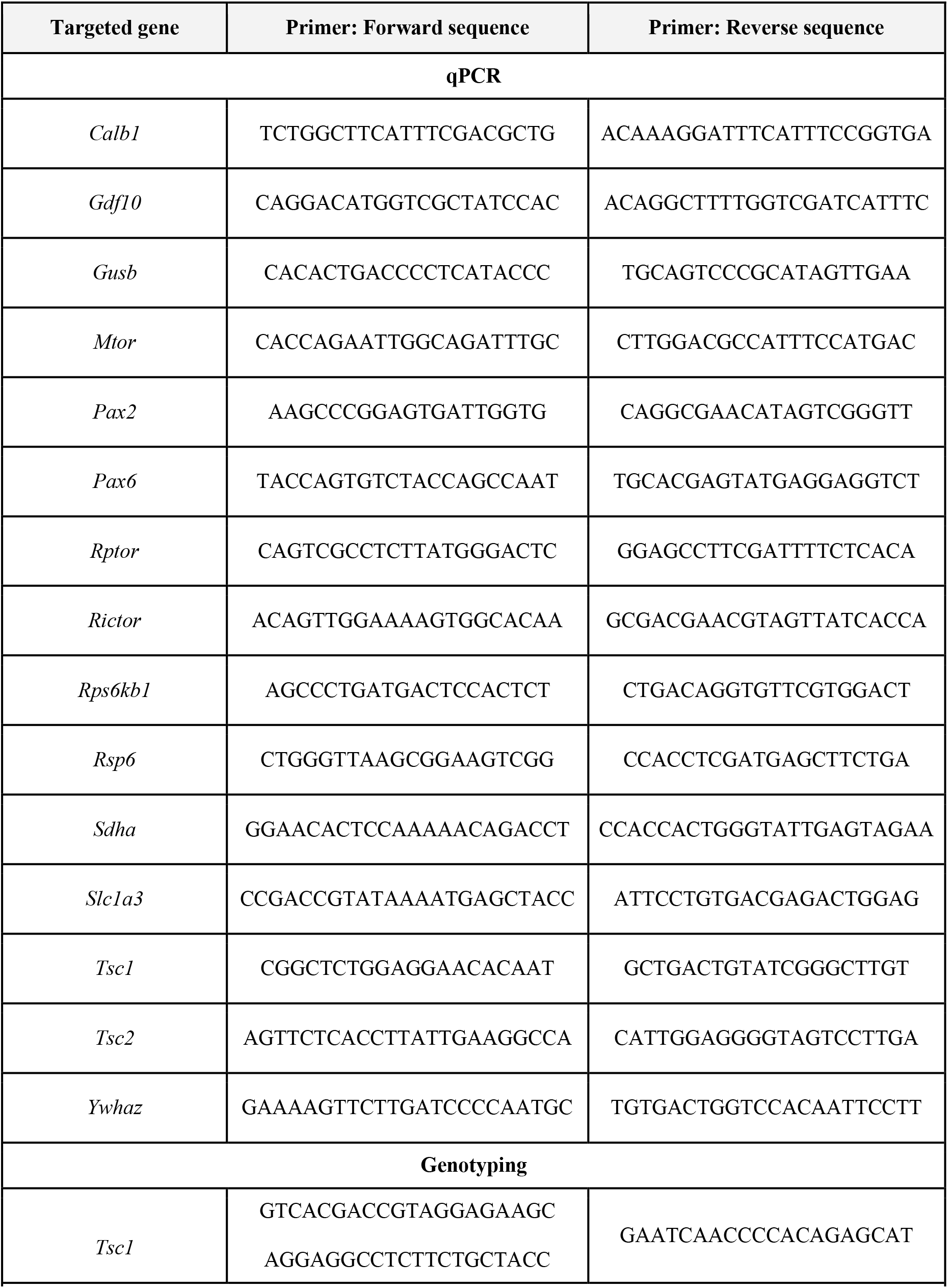
Primer sequences. Overview of genes and primer pairs used for quantitative real time PCR and genotyping of WT and Tsc1^+/-^ mice.

Primers were validated for their specificity *in silico* using *UCSC In-Silico PCR* (genome.ucsc.edu/cgi-bin/hgPcr), *PrimerBank* (pga.mgh.harvard.edu/primerbank) and *blastn* (blast.ncbi.nlm.nih.gov), and *in vitro* with conventional PCR and melt curve analysis.

#### RNA extraction

Following cerebellar dissection, RNA was isolated using a standard chloroform/isopropanol method (39). In brief, tissue in TRI Reagent^®^ (T9424, Sigma) was thawed, homogenized by syringe aspiration (G23 and G25) and vortexed. Chloroform (1:5) was added to the sample, followed by a 5-minute incubation at room temperature (RT). Samples were then centrifuged at 10 800 g, at 4°C for 15 min. The aqueous phase was collected and a 1:1 ratio of isopropanol was added. Samples were centrifuged for 10 min at maximum speed (~20 000 g). The obtained RNA pellet was washed twice with 70% ethanol, air-dried, resuspended in 20 μL of RNase-free water (UltraPure™ DNase/RNase-Free Distilled Water, 10977-035, Invitrogen), and quantified with NanoDrop (Thermo Scientific).

#### RT-qPCR

RNA was transcribed using qScript^®^ cDNA SuperMix (Quantabio, 95048-100), according to manufacturer’s instructions. RT-qPCR was performed with PerfeCTa^®^ SYBR^®^ Green FastMix^®^ (Quantabio, 95072-05K) following manufacturer’s instructions, with 10 μM of forward and reverse primers, and 1 μL cDNA (diluted 1:5). All samples were processed in duplicates. RT-qPCR was performed in a CFX96™ Real-Time PCR detection system (Bio-Rad), with initial denaturation for 1 min at 95°C, followed by 40 cycles of 5 s at 95°C, and 15 s at 55°C, with melting curve generation.

#### Raw data processing

Relative quantification was performed as in (40), on 8 biological samples per genotype for the E15, P1 and P7 groups, and on 6 biological samples for the E18 group. The mean quantitative cycle (Cq) values were extracted for each sample (genes of interest and housekeeping genes), and the mean Cq per gene was calculated within the control group (WT). A ΔCq was then calculated per sample by subtracting the control group average from the sample mean Cq. For each sample, the relative quantities were then calculated ((1+E)^ΔCq^, E=1). Normalized expression per sample (genes of interest) was obtained by dividing the relative quantity of a given sample by the geometric mean of the relative quantities of the two housekeeping genes. The average normalized expression of the samples, in each genotype per gene of interest, was calculated.

### Western blot

Cerebellar tissue from E18 (n = 3 mice per genotype) and P7 (n = 5 mice per genotype), WT and *Tsc1^+/+^* mice, was dissected and immediately frozen in dry ice. Samples were homogenized with a Dounce homogenizer in ice-cold lysis buffer containing 50 mM Tris-HCl pH 8, 150 mM NaCl, 1% Triton X-100, 0.5% sodium deoxycholate, 0.1% SDS and protease inhibitor cocktail (Roche). Protein concentrations were measured using a Pierce BCA protein assay kit (Thermo Fisher). Samples were denatured and proteins separated in SDS-PAGE in Criterion™ TGX Stain-Free™ Gels (Bio-Rad), and transferred onto nitrocellulose membranes with the Trans-Blot^®^ Turbo™ Blotting System (Bio-Rad).

Membranes were blocked with 5% BSA (Sigma-Aldrich) in Tris-buffered saline (TBS)-Tween (20 mM Tris-HCl pH7.5, 150 mM NaCl and 0.1%, Tween20) for 1h, and probed with the following primary antibodies: Pax2 (1:1000, rabbit, Cell Signaling 9666), Phospho-S6 Ribosomal Protein (Ser235/236) (1:1000, rabbit, Cell Signaling 2211), Ribosomal Protein S6 (1:1000, mouse, Santa Cruz SC-74459) or GAPDH (1:1000, mouse, Cell Signaling 97166). Secondary antibodies used were goat anti-Rabbit Immunoglobulins/HRP (1:10000, Agilent Dako P0448) or goat anti-Mouse Immunoglobulins/HRP (1:10000, Agilent Dako P0447). Proteins were detected by the luminol-based enhanced chemiluminescence method (SuperSignal™ West Femto Maximum Sensitivity Substrate or SuperSignal™ West Dura Extended Duration Substrate, Thermo Fisher). Membranes were stripped with Restore™ PLUS Western Blot Stripping Buffer (Thermo Fisher). Densitometry of protein bands of interest was normalised to that of GAPDH using the Image Studio Lite software (LI-COR Biosciences).

### Immunohistochemistry

After collection, E18 embryos (n = 3 mice per genotype) were fixed by immersion in cold 4% paraformaldehyde (PFA) in phosphate buffered saline (PBS). P7 pups (n = 3 per genotype) were injected with an overdose of pentobarbital and transcardially perfused with 4% PFA in PBS. Afterwards, tissue was placed in 4% PFA for 2 hours and transferred into 30% sucrose in 0.1 M phosphate buffer (PB) until embedding. Samples were embedded in 14% gelatine / 30% sucrose and incubated in a 10% PFA / 30% sucrose solution for 1.5 h, at RT, on a shaker. Embedded samples were kept in 30% sucrose / 0.1M PB at 4°C until cut.

Cerebellar samples were cut in 30μm sagittal sections using a cryomicrotome (Leica SM 200R). Free-floating sections were rinsed with PBS and preincubated with 10% normal horse serum (NHS) / 0.5% Triton™ X-100 in PBS, for 1h at RT on a shaker. Sections were then incubated overnight, at 4°C, in 2% NHS / 0.4% Triton™ X-100 in PBS with primary antibodies against Pax2 (1:500, rabbit, Invitrogen 71-6000), Calbindin D28-K (1:10.000, mouse, Sigma C9848) or Parvalbumin (1:500, mouse, Swant 235). The following day, sections were rinsed with PBS and incubated with AlexaFluor 594 (1:500, Donkey anti-rabbit, Jackson 711-585-152) and AlexaFluor 488 (1:500, Donkey anti-mouse, Jackson 715-545-150) in 2% NHS / 0.4% Triton™ X-100 in PBS, for 1.5 h at RT. Sections were rinsed, counterstained with DAPI (1:10.000), and rinsed again with PB before mounting. Sections were imaged with a 10X (E18) or 20X (P7) objective using a Zeiss AxioImager.M2 microscope.

### Microscopy images quantification

The number and area of Pax2^+^ positive cells was automatically counted with Fiji ImageJ (41), using custom-written macros (https://github.com/BaduraLab). Given the positive correlation between nuclear and cell body size, we used the area of Pax2^+^ staining as a proxy for cell area (42). To calculate the distance between Pax2^+^ particles, we used the ND ImageJ plugin (43).

Calbindin-stained sections from P7 WT and *Tsc1^+/-^* mice were used to measure the area of Purkinje cells. The area of 10 randomly selected Purkinje cells per mouse, located between lobules V and VI, was manually measured with Fiji ImageJ, by drawing a region of interest around the visible cell body cross-section.

At P7, parvalbumin (PV) staining was still sparse and dispersed in the developing cerebellum. Thus, we opted for the measurement of the total area occupied by PV stain rather than counting individual cells. To do this, PV-stained cerebellar sections were automatically thresholded and a region of interest was defined including the whole cerebellar section while excluding the already existing Purkinje cell layer (see *Results*). This enabled the measurement of the PV-signal primarily derived from developing molecular layer interneurons (44).

### Statistics

All statistical analysis was performed on GraphPad Prism 8. Data were first screened for the presence of outliers using the ROUT method, and tested for normality using the Shapiro-Wilk test, when applicable. When the normality assumption was followed, a two-tailed t-test was used for data comparison. When this assumption was violated, a two-tailed Mann-Whitney test was used. Variable correlation was performed using Pearson’s correlation, and simple linear regression was used for line fitting.

## Results

### mTOR pathway and cerebellar cell type-specific gene transcription is dysregulated in *Tsc1^+/-^* cerebella

To first investigate whether *Tsc1* haploinsufficiency changes the expression of mTOR pathway and cerebellar genes in the developing cerebellum, we performed RT-qPCR in embryonic and early postnatal cerebellar tissue. Genetic transcription varies greatly during development, hindering the identification of housekeeping genes that remain stable across distinct developmental periods (36). Thus, based on available literature and inter-plate stability, we selected different housekeeping gene pairs for each time point analysed: *Sdha* and *Ywhaz* for E15, *Gusb* and *Sdha* for E18, and *Gusb* and *Ywhaz* for P1 and P7. Within each developmental time point, relative gene expression across the two genotypes, WT and *Tsc1*^+/-^, was compared.

Regarding the transcription of mTOR pathway genes, we found no differences between genotypes at E15, except for the expected down-regulation of *Tsc1* transcription (*t* (13) = 8.685, *p* <0.000001). At E18, in addition to *Tsc1* (*t* (10) = 9.320, *p* = 0.000003), also the transcription of *Tsc2* (*t* (10) = 2.389, *p* = 0.038), *Rictor* (*t* (9) = 2.521, *p* = 0.033) and *Mtor* (*t* (10) = 3.265, *p* = 0.009) was down-regulated in *Tsc1^+/-^* cerebella, suggesting the presence of down-regulation mechanisms regarding mTOR complex genes. At P1, *Tsc1* (*t* (14) = 5.974, *p* = 0.000034) and *Rictor* (*t* (14) = 2.171, *p* = 0.048) were still down-regulated while, at P7, only *Tsc1* (*t* (14) = 3.873, *p* = 0.002) and *Tsc2* (*t* (13) = 2.218, *p* = 0.045) were different from controls (**Figure 1A**).

**Figure 1:**
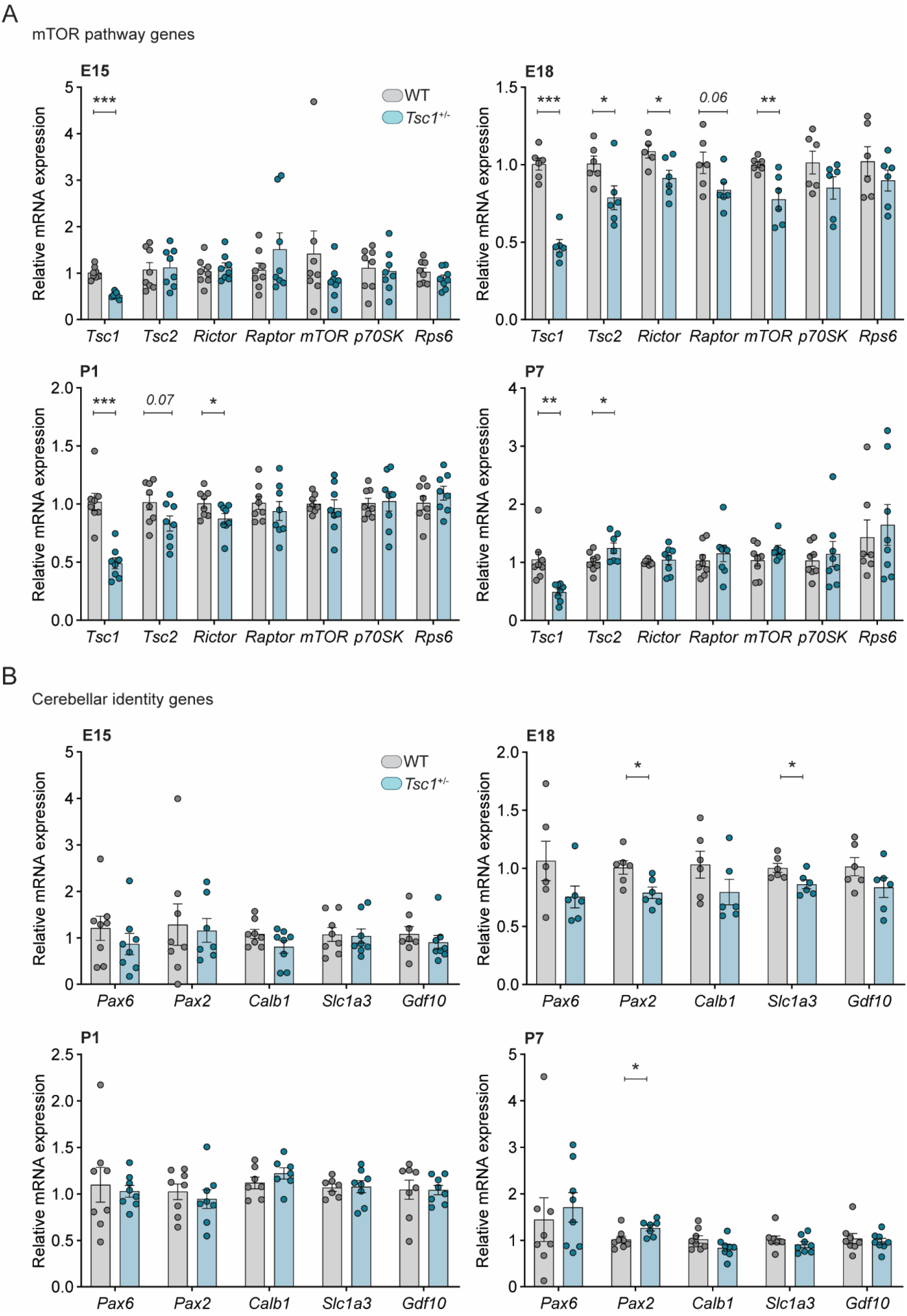
Relative gene expression is altered in *Tsc1^+/-^* cerebella. mRNA expression of mTOR pathway genes (**A**) and cerebellar cell type-specific genes (**B**), relative to housekeeping genes. (**A**) A switch in mTOR-related gene expression occurs at E18, a time point when most markers analysed were significantly down-regulated in *Tsc1^+/-^* cerebella. (**B**) Gene expression of cerebellar identity genes is relatively stable between genotypes, with the exception of *Pax2* and *Slc1a3*. *t*-test, * *p* < 0.05, ** *p* < 0.01, *** *p* < 0.001; n = 8 mice per genotype, except for E18, where n = 6.

We then analysed the transcription of genes involved in the specification of distinct cerebellar cell types. No difference between genotypes was detected at E15 nor at P1. However, at E18, we found that both *Pax2* (*t* (10) = 2.867, *p* = 0.017), a marker for developing interneurons, and *Slc1a3* (*t* (10) = 2.561, *p* = 0.028), a marker for Bergman glia, were down-regulated in *Tsc1*^+/-^ cerebella, while an up-regulation of *Pax2* (*t* (13) = 2.642, *p* = 0.020) was detected in P7 *Tsc1*^+/-^ cerebellar samples (**Figure 1B**). These data indicate that, although the mTOR pathway primarily undergoes post-translational regulation, during development, *Tsc1*^+/-^ haploinsufficiency dysregulates a number of mTOR pathway-related genes, as well as the relative expression of *Slc1a3* and *Pax2*.

### *Tsc1* haploinsufficiency leads to dysregulated gene interactions

Given that E18 and P7 presented with the largest differences between WT and *Tsc1^+/-^* cerebellar gene expression, we focused on these two time points to evaluate the correlation between the relative expression of genes of interest. As expected, we found a significant positive correlation between the relative expression of *Tsc1* and *Tsc2* in WT cerebella (E18: *r*^2^ = 0.65, *p* = 0.052; P7: *r*^2^ = 0.50, *p* = 0.050). While this correlation was still present in *Tsc1^+/-^* mice at E18 (*r*^2^ = 0.91, *p* = 0.003), by P7 this relation was lost in mutants (*r*^2^ = 0.03, *p* = 0.718) (**Figure 2A**). Additionally, while the relative expression of *Tsc1* in WT was negatively correlated with the relative expression of *S6* (E18: *r*^2^ = 0.83, *p* = 0.011; P7: *r*^2^ = 0.64, *p* = 0.03), this was not the case for *Tsc1^+/-^* mice (E18: *r*^2^ = 0.01, *p* = 0.82; P7: *r*^2^ = 0.03, *p* = 0.71) (**Figure 2B**). This indicates that genetic mTOR pathway dysregulation in *Tsc1^+/-^* cerebella can be found early in development, likely prior to detectable protein changes, and that these deficits exhibit time-dependent progression.

**Figure 2:**
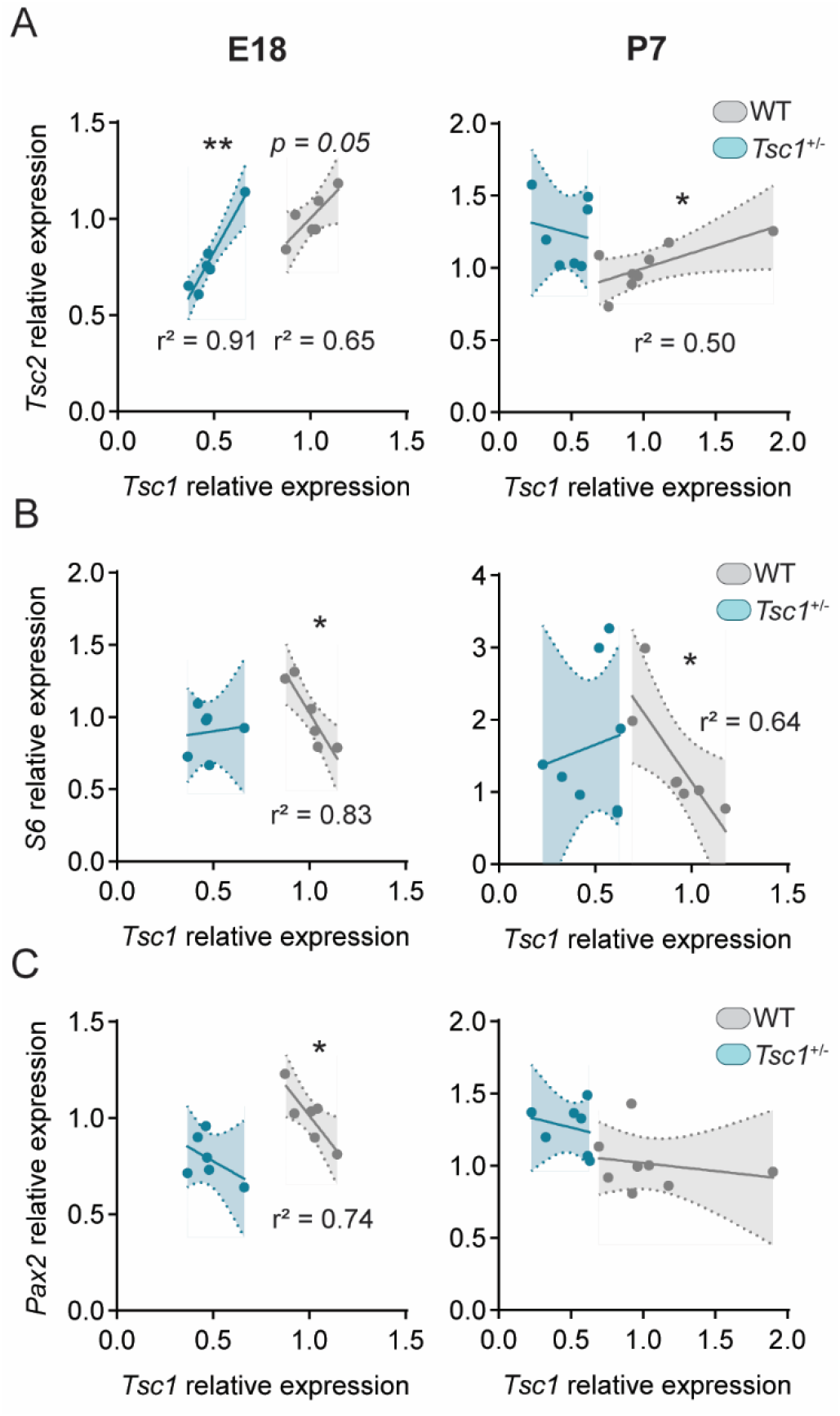
Gene correlation is lost in *Tsc1^+/-^* cerebella. Pearson’s correlation plots for mRNA relative expression of *Tsc1* over (**A**) *Tsc2*, (**B**) *S6* and (**C**) *Pax2* in E18 (left column) and P7 (right column) cerebella. Pearson’s correlation, * *p* < 0.05, ** *p* < 0.01; n = 8 mice per genotype, except for E18, where n = 6.

Pax2 expression is initiated in inhibitory interneuronal precursors during their last mitosis (45). Possibly reflecting the positive role of the mTOR pathway on neuronal differentiation (46), we found a negative correlation between *Tsc1* and *Pax2* relative expression in E18 WT cerebella (*r*^2^ = 0.74, *p* = 0.03). However, this correlation was absent in *Tsc1^+/-^* mice (*r*^2^ = 0.22, *p* = 0.35) (**Figure 2C**). This suggests that mTOR pathway disruption through *Tsc1* haploinsufficiency could lead to early dysfunction of *Pax2^+^* cell differentiation.

### *Tsc1* haploinsufficiency increases the size of Pax2^+^ cells but does not affect cell number at E18

The expression of *Pax2* in the cerebellum is initiated at around E13.5, primarily found in Golgi cells, and continues into postnatal time points, with the differentiation of stellate and basket cells perinatally (47–49). Having found that the relative mRNA expression of *Pax2* was down-regulated in *Tsc1^+/-^* cerebella at E18, we then investigated the protein expression of Pax2 at this time point. Using western blot in whole cerebellar extracts, we detected no differences between genotypes regarding the expression of Pax2 (*U* = 4, *p* > 0.99) (**Figure 3A**). Correspondingly, we also found that WT and *Tsc1^+/-^* cerebella presented with a similar number of Pax2-positive (Pax2^+^) cells (*U* = 127.5, *p* = 0.16) (**Figure 3B, C**).

**Figure 3:**
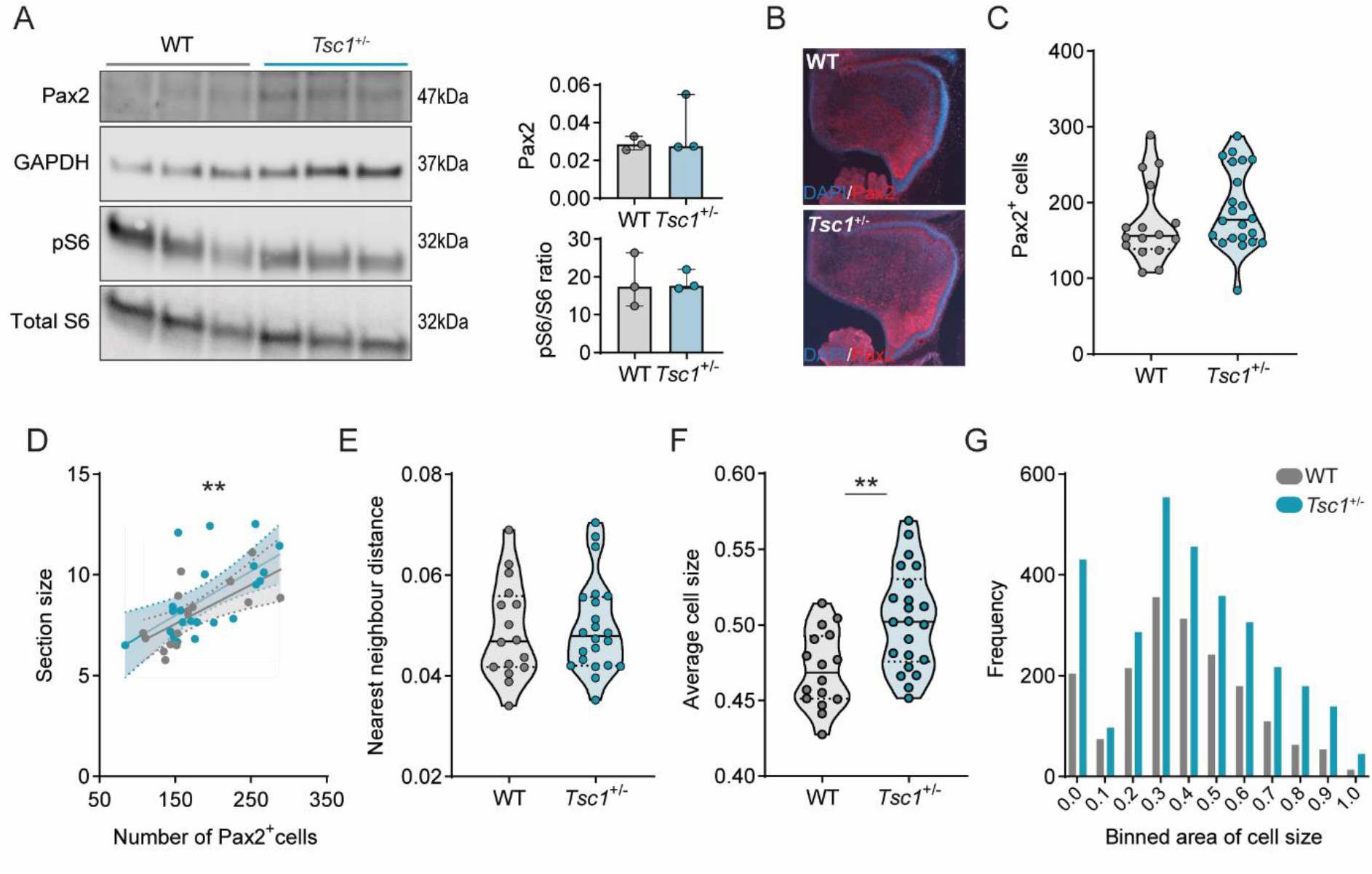
Increased Pax2^+^ cell size is found in E18 *Tsc1^+/-^* cerebella. (**A**) Western blot quantification of Pax2, phosphorylated S6 (Ser235/236) and total S6, in whole cerebellar tissue (n = 3 mice per genotype, Mann-Whitney test). (**B**) Representative Pax2-stained sagittal section (red) counterstained with DAPI (blue). (**C**) Pax2^+^ cell count on WT (n = 16 sections from 2 mice) and *Tsc1^+/-^* (n = 22 sections from 2 mice) sagittal sections (Mann-Whitney test). (**D**) Pearson’s correlation between cerebellar section size and the number of Pax2^+^ cells (WT in grey, *Tsc1^+/-^* in blue; 16 points for WT and 22 for *Tsc1^+/-^* mice). (**E-F**) Nearest neighbour distance and average cell size of WT and *Tsc1^+/-^* Pax2^+^ cells (n = 16 sections from 2 mice for WT and n = 22 sections from 2 mice for *Tsc1^+/-^*; *t*-test). (**G**) Frequency of Pax2^+^ cells over the binned cell area (2863 cells from WT and 4252 cells from *Tsc1^+/-^* mice). Area in inches. ** *p* < 0.01.

At this stage, Pax2^+^ cells are highly migratory and quasi-uniformly dispersed through the developing cerebellum (45). We corroborated this uniform positioning in both WT and *Tsc1^+/-^* sections as, in the two genotypes, the total number of Pax2^+^ cells was positively correlated with the correspondent section size (WT: *r*^2^ = 0.44, *p* = 0.005; *Tsc1^+/-^*: *r*^2^ = 0.36, *p* = 0.003) (**Figure 3D**). To investigate the position of these cells, we then calculated the nearest neighbour distance between Pax2^+^ cells, and used this measure as a proxy for cell migration. We found that both WT and *Tsc1^+/-^* Pax2^+^ cells were separated by similar distances (*t* (36) = 0.26, *p* = 0.84) (**Figure 3E**). This suggests that, while *Tsc1* haploinsufficiency causes a reduction in *Pax2* transcription, this is not sufficient to affect the generation nor migration of Pax2^+^ cells at this stage in development.

Loss of function of *Tsc1*, and consequent mTOR overactivation, is often accompanied by changes in neuronal cell size (50). Thus, we then measured the size of Pax2^+^ cells in the E18 cerebellum. Despite not finding increased overall levels of pS6 (Ser235/236)/total S6 in whole cerebellar extracts of *Tsc1^+/-^* mice (*U* = 4, *p* > 0.99) (**Figure 3A**), we found that these mice presented with enlarged Pax2^+^ cells when compared to WT cells (*t* (36) = 3.38, *p* = 0.002) (**Figure 3F**). These larger cells were present across the full spectrum of measured areas (**Figure 3G**). This data suggests that the mTOR pathway is overactive in *Tsc1*^+/-^ interneuronal progenitor cells, leading to an overall increase in cell size that seems to affect distinct lineages of inhibitory interneurons.

### Overactive mTOR pathway in P7 cerebellum leads to perturbed interneuron development in *Tsc1^+/-^* mice

At P7, although the vast majority of cerebellar interneurons is already generated, these can be found in distinct developmental stages, from migrating to fully mature neurons (49,51). To further explore the increase in *Pax2* relative expression we previously found in *Tsc1*^+/-^ cerebella at P7, we next analysed the expression and distribution of Pax2^+^ cells in the P7 cerebellum. As observed for the E18 cohort, we found no difference in Pax2 protein expression between WT and *Tsc1^+/-^* cerebella (*t* (8) = 1.61, *p* = 0.76) (**Figure 4A**). Supporting our qPCR data, immunohistochemical analysis of Pax2-labelled cerebellar sections revealed that *Tsc1^+/-^* cerebella presented with an increased number Pax2^+^ cells (**Figure 4B, C**). In addition, these cells were most abundant in the cerebellar white matter (**Figure 4B**), altogether suggesting the presence of an immature Pax2 phenotype in *Tsc1^+/-^* mice (52,53).

**Figure 4:**
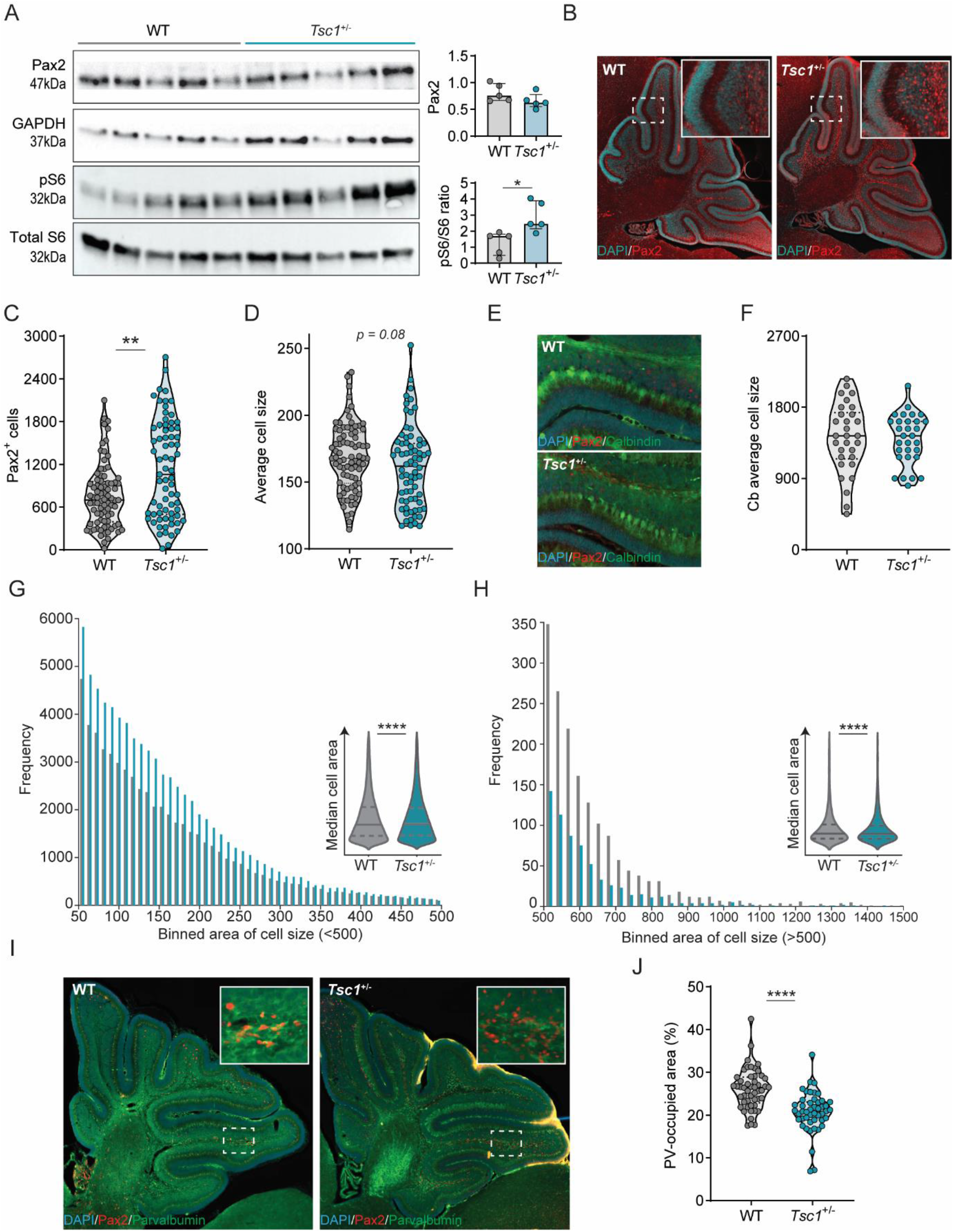
Tsc1^+/-^ mice present with interneuron development and maturation deficits at P7. (**A**) Western blot quantification of Pax2, phosphorylated S6 (Ser235/236) and total S6, in whole cerebellar tissue (n = 5 mice per genotype; *t*-test). (**B**) Representative Pax2-stained sagittal section (red) counterstained with DAPI (blue). (**C**) Pax2^+^ cell count on WT (n = 81 sections from 3 mice) and *Tsc1^+/-^* (n = 71 sections from 3 mice) sagittal sections (Mann-Whitney test). (**D**) Average cell size of WT and *Tsc1^+/-^* total Pax2^+^ cells (n = 81 sections from 3 WT mice and n = 71 sections from 3 *Tsc1^+/-^* mice; Mann-Whitney test). (**E**) Representative sagittal section stained for Calbindin (green), Pax2 (red) and counterstained with DAPI (blue). (**F**) Average cell size of WT and *Tsc1^+/-^* Purkinje cells, measured between lobules V and VI (n = 10 cells per mouse, 3 mice per genotype; *t*-test). (**G-H**) Frequency of small (left) and large (right) Pax2^+^ cells over binned cell area (61 189 cells from WT and 79 955 cells from *Tsc1^+/-^*, 3 mice per genotype). Insets show the median area of the measured cells. (**I**) Representative section stained for PV (green), Pax2 (red) and DAPI (blue) (Mann-Whitney test). (**J**) Percentage of PV-occupied area per section (n = 51 section from 3 WT mice and n = 49 sections from 3 *Tsc1^+/-^* mice; Mann-Whitney test). Area in pixels. * *p* < 0.05, ** *p* < 0.01, *** *p* < 0.001.

In contrast with the results obtained at E18, the size of Pax2^+^ cells tended to be smaller in *Tsc1^+/-^* mice compared to WT (*U* = 2329, *p* = 0.08) (**Figure 4D**). This putative Pax2^+^ cell size change appeared to be specific towards Pax2^+^ cells, as we found no difference in the size of other GABAergic cells, such as Purkinje cells (*t* (58) = 0.1219, *p* = 0.90) (**Figure 4E, F**). This was an intriguing finding given that we detected an overall increase in pS6 (Ser235/236) / total S6 in whole cerebellar extracts, suggesting mTOR pathway hyperactivation (**Figure 4A**). Thus, we further investigated Pax2^+^ cell size by analysing their frequency distribution across distinct binned areas. We found that *Tsc1^+/-^* mice presented with an increased number of small Pax2^+^ cells (**Figure 4G**), which likely represent stellate and basket cells in the nascent molecular layer (45). Contrarily, *Tsc1^+/-^* mice exhibited decreased numbers of large Pax2^+^ cells, indicative of the granular layer interneurons, Golgi cells (44,54) (**Figure 4H**). Furthermore, small Pax2^+^ cells in *Tsc1^+/-^* mice were slightly smaller than WT cells (WT = 132 vs *Tsc1^+/-^* = 123, median size, *p* < 0.0001) (**Figure 4G-inset**), as were large the Pax2^+^ cells (WT = 584 vs *Tsc1^+/-^* = 572 median size, *p* < 0.0001) (**Figure 4H-inset**). This suggests that *Tsc1* haploinsufficiency differentially alters Pax2^+^-derived lineages.

### Maturation of interneuron precursors at P7 is altered in *Tsc1^+/-^* cerebella

The density of cerebellar interneurons remains relatively constant between P5 to P10 (55). For the majority of these cells, comprising mostly of the molecular layer interneurons, down-regulation of Pax2 is accompanied by up-regulation of PV, a process that marks interneuronal maturation (44,56). Given that we found an increase in the number of Pax2^+^ cells in *Tsc1^+/-^* cerebella at P7 but not at E18, we speculated that this could reflect a decrease in Pax2 down-regulation rather than an overall increase in cell generation. Therefore, we focused on the maturation of the interneuronal cells.

As a proxy of cell maturation, we quantified the percentage of cerebellar P7 area occupied by PV staining (*see methods*) (**Figure 4I**). We found that *Tsc1^+/-^* cerebellar sections presented with a decreased percentage of surface area labelled by PV staining when compared to WT mice (median percentage: 26.45% for WT vs 20.89% for *Tsc1^+/-^*, *p* < 0.0001), suggesting that *Tsc1* haploinsufficiency is accompanied by deficits in the maturation of molecular layer interneurons (**Figure 4J**).

## Discussion

The mTOR pathway has been linked to many cellular and metabolic events, including cell replication, growth and biomass production (1). Although the mTOR kinase is known to be required for the formation of the central nervous system (CNS) (2,4), its precise role in the development of distinct cell lineages is not completely understood. Currently, a tight regulation of the timing of mTOR pathway activation appears to be essential for the balance between undifferentiated cell proliferation and cell differentiation. In postnatal mice, mTOR signalling is detected in proliferating neural stem cells (NSCs), while knockdown of its activity reduces proliferation. Conversely, increasing mTOR activation leads to a higher number of terminally differentiated NSCs at the expense of their renewal (4). Further, in cortical interneuron progenitors, deletion of mTOR decreases proliferation, leading to a reduction of mature calbindin-positive cells (57).

In the developing cerebellum, few reports have addressed the role of the mTOR pathway in the specification of distinct cell types. Proper mTOR signalling appears imperative for the correct development of Purkinje cells (PC), as disruptions in either mTORC1 or mTORC2 signalling in PC from E17.5 lead to smaller soma size and deficits in dendritic arborization (58,59). Conditional deletion of *Rictor*, essential for mTORC2 kinase activity, from all CNS precursor cells at E10.5, induces early postnatal changes to PC, including the emergence of several primary dendrites and abnormal vermal macrostructure (59). Furthermore, while the loss of *Rictor* does not seem to affect the development of cerebellar granule neuron precursor cells (GNP), deleting *Raptor*, necessary for mTORC1 activity, leads to a decrease in cell number (60). Additionally, increased S6 kinase activity leads to a reduction in proliferating GNP due to premature cell cycle exit (61).

To better understand how global mTOR overactivation impacts embryonic and early postnatal cerebellar development, we used *Tsc1*^+/-^ mice, often used as a mouse model to study mTORopathy-associated ASD. Because the mTOR kinase is an important regulator of translation (1), we first evaluated expression levels of mTOR pathway-related and cerebellar cell-specific genes in the developing cerebellum. The overall loss of correlation in distinct gene pairs we found in *Tsc1^+/-^* mice supports a deficient stability of the TSC1-TSC2 complex, as well as a dysregulation of translational machinery. This is in line with previous work demonstrating that increased mTOR function leads to an altered profile in neuronal genetic transcription (62). Thus, *Tsc1^+/-^* haploinsufficiency alters the translational landscape of the cerebellum early in development, through the dysregulation of central mTOR-sensitive genes.

To identify which cell types could potentially be more susceptible to mTOR overactivation, we analysed the relative expression of cerebellar-specific cell markers in the developing cerebellum. We found cerebellar lineage deficits as early as E18, which were further evident in the first week of postnatal development. Specifically, we found that cerebellar interneuronal precursors, characterized by the expression of *Pax2*, seemed to be particularly sensitive to global haploinsufficiency of *Tsc1* and the consequent mTOR pathway overactivation. We observed that these interneuron precursors presented with hyperactive mTOR pathway. Additionally, this overactivation appeared to differentially affect the development of molecular and granular layer interneurons. The observed changes in *Pax2* mRNA expression indicate that *Tsc1* haploinsufficiency leads to a delay in the initiation of Pax2 expression in embryonic development, causing its increased expression during later postnatal periods. Thus, it is possible that disruption of cerebellar mTOR signalling primarily affects the maturation of interneurons rather than progenitor cell pool maintenance. Alternatively, the deficient down-regulation of mTOR signalling found in P7 *Tsc1^+/-^* mice could also contribute to an increase in cell proliferation, leading to elevated overall numbers of Pax2^+^ cells. These are pertinent hypotheses, as recent work in *Drosophila* has demonstrated direct Pax2 expression modulation by the mTOR pathway (63). In *Drosophila*, D-Pax2 is a main regulator of cell fate in the developing eye (63), and was shown to physically interact with Unkempt (Unk), a highly conserved zinc finger/RING domain protein, which is also highly expressed in mouse cerebellum (64). In *Drosophila*, Unk expression is negatively regulated by the mTOR pathway and *Tsc1* mutant flies present with increased D-Pax2 expression (63).

Using immunocytochemistry, we found deficits in molecular layer interneuron maturation, as evidenced by decreased PV staining in the cerebellum of *Tsc1^+/-^* P7 mice. PV is a calcium binding protein, abundant in Purkinje cells and cerebellar molecular layer interneurons (65). Thus, we focused on this population as they make up the majority of cerebellar interneuronal cells. During brain development, the initiation of PV expression coincides with the expression of a number of synaptogenesis markers, such as solute carrier family 32 and GABAA receptor α1 subunit (44). Furthermore, the amount of PV expression was shown to determine presynaptic calcium dynamics in cerebellar interneurons, modulating neurotransmitter release (44). Thus, the changes in PV expression levels and timing of the interneuronal maturation that we found in *Tsc1^+/-^* mice, could potentially lead to deficits in synaptic integration in the cerebellum (66).

Behaviourally, *Tsc1* mice models present with ASD-like features, including decreased social interaction, increased repetitive behaviours and deficient reversal learning (29,31). This is a similar phenotype to the one found in PV knockout mice (67). Conversely, decreased numbers of PV positive cells are found in other models of ASD, namely *Cntnap2^-/-^, Shank1^-/-^, Shank3B^-/-^*, and *Brinp3^-/-^* (68–71). Based on this evidence, a recent review by Filice and colleagues proposed the “Parvalbumin Hypothesis of Autism Spectrum Disorder”, in which down-regulation of parvalbumin expression leads to altered neuronal function and abnormal neurotransmitter release, in addition to increasing reactive oxygen species production and dendritic branching (72). Thus, deficits in PV could be one of the mechanisms integrating distinct high-risk mutations that lead to the development of ASD.

## Acknowledgments

We thank Ype Elgersma for the total S6 antibody; Chris de Zeeuw for the Pax2 and S6 antibodies; Roxanne ter Haar for the help in mouse genotyping and breeding; Elize Haasdijk and Ivy Hau for their help with immunohistochemistry tissue processing. This research was supported by the Netherlands Organization for Scientific Research and ZonMw (VIDI/917.18.380,2018; AB).

## Author contributions

IS, AS, and AB designed the study and analysis. IS, AS and MRO performed the qPCR and histological experiments. CO executed the western blot experiments. IS, AS, MRO and CO analysed the data. CT, MS and AB supervised the project. IS, AS and AB wrote the first draft. All authors edited the manuscript.

## Data and code availability

The raw data that support the findings of this study is available from the corresponding author upon request. The code is deposited at https://github.com/BaduraLab.

